# The effects of resistance training on denervated myofibers, senescent cells, and associated protein markers in middle-aged adults

**DOI:** 10.1101/2023.10.04.560958

**Authors:** Bradley A. Ruple, Madison L. Mattingly, Joshua S. Godwin, Mason C. McIntosh, Nicholas J. Kontos, Anthony Agyin-Birikorang, J. Max Michel, Daniel L. Plotkin, Shao-Yung Chen, Tim. N. Ziegenfuss, Andrew D. Fruge, L. Bruce Gladden, Austin T. Robinson, C. Brooks Mobley, Abigail L. Mackey, Michael D. Roberts

## Abstract

Denervated myofibers and senescent cells are hallmarks of skeletal muscle aging. However, sparse research has examined how resistance training affects these outcomes. We investigated the effects of unilateral leg extensor resistance training on denervated myofibers, senescent cells, and associated protein markers in middle-aged participants (MA, 55±8 years old, 17 females, 9 males). We obtained vastus lateralis (VL) muscle cross-sectional area (mCSA), VL biopsies, and strength assessments before and after training. Fiber cross-sectional area (fCSA), satellite cells (Pax7+), denervated myofibers (NCAM+), senescent cells (p16+ or p21+), senescence-related proteins, and senescence-associated secretory phenotype (SASP) proteins were analyzed from biopsied muscle. Leg extensor peak torque increased after training (p<0.001), while VL mCSA trended upward (p=0.082). No significant changes were observed for fCSA, NCAM+ myofibers, or senescent (p16+ or p21+) cells, albeit satellite cells increased after training (p=0.037). While >90% satellite cells were not p16+ or p21+, most p16+ and p21+ cells were Pax7+ (>90% on average). Training altered 13/46 proteins related to muscle-nerve communication (all upregulated, p<0.05) and 10/19 proteins related to cellular senescence (9 upregulated, p<0.05). Only 1/17 SASP proteins increased with training (IGFBP-3, p=0.031). In conclusion, resistance training upregulates proteins associated with muscle-nerve communication in MA participants but does not alter NCAM+ myofibers. Moreover, while training increases senescence-related proteins in skeletal muscle, this coincided with an increase in satellite cells but not alterations in senescent cell content or SASP proteins. Hence, we interpret these collective findings as resistance training being an unlikely inducer of cellular senescence in humans.

## INTRODUCTION

Skeletal muscle is a tissue that is responsible for force generation, locomotion, metabolism, and heat production [1]. Skeletal muscle consists of thousands of muscle fibers, or myofibers, and each fiber is innervated by a single alpha motor neuron branch. This neuron, and all the myofibers it innervates, is collectively termed a motor unit. Functionally, the motor unit is responsible for relaying signals from the nervous system to the muscle to elicit muscle contraction. However, the site of muscle-nerve communication can be altered, specifically the neuromuscular junction (NMJ) [2, 3]. When a motor neuron is no longer communicating with a myofiber, this is referred to as denervation [4]. With aging there is an acceleration of denervation and a concomitant decline in reinnervation [5, 6]. Additionally, a preferential denervation of type II fibers occurs whereby the denervated myofibers are re-innervated by lower-threshold motor units, a phenomenon termed fiber type grouping [3]. Another well-established characteristic of aging is age-related muscle loss [7], and losses in myofiber innervation could partially explain why age-related atrophy ensues. In this regard, neural cell adhesion molecule positive (NCAM+) myofibers (indicative of denervation) are higher in older versus younger individuals, thus providing evidence that this phenomenon occurs [8, 9]. While there have been multiple studies showing the benefits of exercise for older individuals [10-14], there is limited and equivocal research on how exercise training affects denervated myofibers [15-18]. Furthermore, older individuals exhibit blunted hypertrophic responses to resistance training, and a recent meta-analysis supports that increased myofiber hypertrophy to resistance training is inversely related with age [19]. These data could be explained, in part, by the presence of more denervated myofibers and/or altered NMJ characteristics in aged individuals.

A diminished hypertrophic response to resistance training in older individuals may also be, in part, due to an increased presence of senescent cells. p16 (cyclin dependent kinase inhibitor 2A; p16^INK4a^) and p21 (cyclin dependent kinase inhibitor 1A; p21^WAF1/CIP1^) are markers of senescence that inhibit cyclin-dependent kinases and promote cell cycle arrest [20-22]. Senescent cells are defined as a cell type in the interstitium that enters a state of semi-permanent cell cycle arrest and loses its regenerative capacity, while secreting molecules associated with inflammation (i.e., senescence-associated secretory phenotype, or SASP) [23]. Aging is positively associated with senescent cell abundance in muscle tissue [24-26]. Specifically, older mice present more senescent cells in the extracellular matrix of the plantaris muscle in response to synergist ablation-induced mechanical overload (but not in the basal state) relative to younger adult mice and this coincides with an impairment in muscle hypertrophy [27]. However, a senolytic cocktail (i.e., dasatinib and quercetin, or D+Q) reduced the presence of senescent cells in older mice and rescued the impaired hypertrophic phenotype. While these data are informative, little is known regarding the association between senescent cell abundance and the hypertrophic response to resistance training in older humans.

Given the knowledge gaps in humans discussed above, the purpose of this study was multifaceted. First, we sought to examine how eight weeks of unilateral knee extensor resistance training affected the number of denervated (NCAM+) myofibers and senescent cells (p16+ or p21+) in middle-aged individuals (MA) in both the trained and non-trained legs. Additionally, we implemented deep proteomics and targeted antibody array approaches to examine how proteins associated with cellular senescence and muscle-nerve communication were affected in the trained leg of these individuals. As with some of our past research [28, 29], we also sought to examine if resistance training elicited rejuvenating effects on assayed targets. Thus, tissue from a college-aged cohort (denoted as “Y” throughout) was used to examine if any aging or “training-rejuvenating” effects existed for the outcome variables. We hypothesized that, prior to training, both legs of the MA participants would possess more denervated myofibers and senescent cells compared to Y participants. We also hypothesized that resistance training would decrease the number of denervated myofibers and associated proteins in the training leg. Finally, given that senescent cells increase with age and stress, we hypothesized that senescent cells and proteins associated with cellular senescence would increase in the trained leg of MA participants.

## METHODS

### Ethical approval and participant screening

This study was a non-related sub-aim of a study approved by the Auburn University Institutional Review Board (IRB protocol #21-461 MR 2110) which aimed to investigate the effects of a nutritional supplement (312 mg of combined Wasabia japonica extract, theacrine, and copper (I) niacin chelate) versus a placebo on unrelated markers over an eight-week period. A unilateral leg resistance training (2d/week) protocol was implemented to perform non-supplementation secondary analyses as presented herein. To ensure supplementation did not have appreciable effects on the assayed targets in skeletal muscle, we statistically compared the placebo and supplementation groups for pre-to-post intervention change scores in phenotypes (n=13 per group) histology (n=10 per group), targeted proteomics (n=9 per group), and SASP protein markers (n=6-7 per group). No significant differences in these scores were observed between supplementation groups for the following variables in the training leg or non-training leg: vastus lateralis (VL) muscle cross-sectional area (mCSA) (p=0.693 and p=0.919, respectively), mixed fiber cross-sectional area (fCSA) (p=0.187 and p=0.333, respectively), the percentage of NCAM+ myofibers (p=0.803 and p=0.768, respectively), satellite cell number (p=0.131 and p=0.705, respectively), or p21+/Pax7+ cell counts (p=0.360 and p=0.322, respectively), or p16+/Pax7+ cell counts (p=0.376 and p=0.900, respectively). Of the 66 targeted proteins from the proteomic analysis (only performed in the trained leg), only one was affected with supplementation (MTR, related to “axon regeneration”, was downregulated, p=0.043). Of the 17 targeted SASP proteins (only performed in the trained leg), five showed significant change score differences between supplementation groups exceeding p<0.05 according to the Mann-Whitney U statistic (GM-CSF, IGFBP3, IL-6, IL-8, and RANTES). Wilcoxon rank tests indicated that the increase in IGFBP3 was the only significant SASP marker to increase in the 6 participants that consumed the nutritional supplement (p=0.031). Conversely, Wilcoxon rank tests indicated that IL-6 (p=0.047) and IL-8 (p=0.016) significantly decreased in the 7 participants that consumed the placebo supplement. Hence, given the marginal differences between supplementation groups for training outcomes, histology markers, proteomic targets, and SASP proteins, we opted to combine the two groups to robustly increase our statistical power for this secondary analysis.

Tissue from two prior studies were also used (Protocol #19–249 MR 1907 and #20–136 MR 2004) whereby strength, VL ultrasound images, and VL biopsies were obtained from younger adults [30, 31]. While the younger participants were involved in training studies, the participants’ basal (pre-intervention) characteristics were used to examine whether training leg outcomes in MA participants were altered to “youth-like” levels. Due to tissue or resource limitations, subsets of MA and Y participants were analyzed for certain assays, and these details are provided in Table 1.

**Table 1.** Participant n-sizes for outcome variables.

| Outcome | MA participants | Y participants |
| --- | --- | --- |
| General phenotypes | 26 (age: 55±8 years old; 9M/17F) | 15 (age: 22±3 years old; 6M/9F) |
| Histology variables | 20 (age: 55±8 years old; 5M/15F) | 11 (age: 22±3 years old; 5M/6F) |
| Targeted proteomics* | 18 (age: 55±8 years old; 4M/14F) | 9 (age: 22±2 years old; 2M/7F) |
| SASP proteomics* | 13 (age: 58±8 years old; 3M/10F) | 5 (age: 22±1 years old; 1M/4F) |

Notes: \*, indicates that assays were only performed in the trained leg of MA participants.

Inclusion criteria for all studies indicated that participants had to possess a body mass index <34 kg/m^2^, have no or minimal experience with resistance training (≤ 1 day/week) one year prior to the study, and be free of any medical condition that would contraindicate participating in an exercise program or donating skeletal muscle biopsies. Participants provided verbal and written consent to participate prior to data collection and the study conformed to standards set by the latest revision of the Declaration of Helsinki except for being registered as a clinical trial.

### Study Design and Resistance Training Paradigm

The resistance training intervention consisted of supervised unilateral leg extensions (2d/week for 8 weeks), and the intervention was preceded and followed by strength and VL muscle assessments (described later). For logistical purposes, all MA participants trained their right legs, with the left leg serving as a within-participant, untrained control leg. From 3-repetition max (3RM) testing (described below), 13 of the 26 right legs were considered the dominant leg.

For each training day, participants performed five sets of 12 repetitions per session. The beginning training load was established at ∼40% of the participant’s 3RM (described below). After each set, participants verbally articulated their perceived repetitions in reserve (RIR), and training load was adjusted accordingly. RIR values of 0-2 after a set resulted in no training load change in each session. RIR values of 3-5 for consecutive sets resulted in the training load being increased by 5-10%. For RIR values ≥6 after one set, the training load was increased by 10-20%. If the weight could not be performed with full range of motion, or the participant could not complete 12 repetitions for a given set, the training load was decreased.

### Strength testing

The first and last workout of the eight-week training paradigm consisted of maximal leg extensor-flexion torque assessments using isokinetic dynamometry (Biodex System 4; Biodex Medical Systems, Inc., Shirley, NY, USA) and 3RM leg extensor strength testing. Notably, both legs were used for these assessments. Prior to dynamometer testing, the participant’s lateral epicondyle was aligned with the axis of the dynamometer’s lever arm, and the hip was positioned at 90°. The starting leg was randomized for everyone prior to each test. The participant’s shoulders, hips, and leg were strapped and secured for isolation during testing. Following three warm-up trials at a submaximal effort, participants completed five maximal voluntary isokinetic knee extension and flexion actions at 60 degrees/second. Participants were provided verbal encouragement during each contraction. The isokinetic contraction resulting in the greatest peak torque value was used for analyses. Participants were given a one-minute rest and then the contralateral leg completed this exact protocol. Approximately five minutes following isokinetic dynamometry testing, participants performed 3RM strength testing on both the right and left leg. Prior to testing, participants were given a warm-up load and instructed to complete 10 repetitions. After participants recorded their RIR for the warmup set, the weight was adjusted accordingly for another warm-up set of five repetitions. RIR was recorded again to determine the participant’s starting load for a 3RM attempt. The load was incrementally increased 5-10% per 3RM attempt until 3RM testing concluded, indicated by failure of full range of motion on any of the repetitions, or if RIR recorded was 0. Participants were allowed a full three minutes of recovery between attempts. The isokinetic dynamometry and 3RM testing described herein was similar for both the first and final workout.

### Testing Sessions

#### Urine specific gravity testing for hydration

Participants performed a testing battery prior to the start of training (PRE) and 3-5 days following the last resistance training workout (POST). Participants arrived for testing at a minimum of 4 hours fasted and well hydrated. Upon arrival participants submitted a urine sample (∼5 mL) for urine specific gravity assessment (USG). Measurements were performed using a handheld refractometer (ATAGO; Bellevue, WA, USA). USG levels in all participants were ≤ 1.020, indicating sufficient hydration [32].

#### Body composition testing

Once adequate hydration was determined, body composition was assessed using multi-frequency bioelectrical impedance analysis (InBody 520, Biospace, Inc. Seoul, Korea). From the scan, body fat percentage was recorded. Previously determined test-retest reliability yielded an intraclass correlation coefficient (ICC_3,1_) of 0.99, standard error of the measurement (SEM) of 0.87%, and minimal difference (MD) of 1.71% for body fat percentage.

#### Ultrasonography assessment for muscle morphology

A detailed description of VL assessments using ultrasonography has been published previously by Ruple et al. [33]. Briefly, real-time B-mode ultrasonography (NextGen LOGIQe R8, GE Healthcare; Chicago, IL, USA) using a multifrequency linear-array transducer (L4-12T, 4–12 MHz, GE Healthcare) was used to capture right and left leg images to determine VL muscle cross-sectional area (mCSA). Prior to scans, the mid-thigh of the leg was determined by measuring the total distance from the mid-inguinal crease in a straight line to the proximal patella, with the knee and hip flexed at 90°, a mark was made using a permanent marker at 50% of the total length. From that location, a permanent marker was used transversely to mark the mid-belly of the VL. This marking is where all PRE ultrasound images were taken as well as the muscle biopsy (described below). POST images were taken at the PRE biopsy scar to ensure location consistency between scans. During mCSA scans, a flexible, semirigid pad was placed around the thigh and secured with an adjustable strap to allow the probe to move in the transverse plane. Using the panoramic function of the device (LogicView, GE Healthcare), images were captured starting at the lateral aspect of the VL and moving medially until rectus femoris was visualized, crossing the marked location. All ultrasound settings were held constant across participants and laboratory visits (frequency: 10 MHz, gain: 50 dB, dynamic range: 75), and scan depth was noted and held constant across time points per participant. Images were downloaded and analyzed offline using ImageJ software (National Institutes of Health, Bethesda, MD). All ultrasound images were captured and analyzed by the same investigators at each timepoint. Previously determined test-retest reliability on 10 participants measured twice within 24 h (where BAR captured images and JSG analyzed images) yielded an ICC of 0.99, SEM of 0.60 cm^2^, and MD of 1.65 cm^2^ for VL mCSA.

#### Collection of muscle tissue

Muscle biopsies were obtained at PRE and POST, from the mid-belly of both the right and left VL muscle, with sampling time of day at PRE and POST being standardized for each participant. Lidocaine (1%, 1.0 mL) was injected subcutaneously superficial to the skeletal muscle fascia at the previously marked location. After five minutes of allowing the anesthetic to take effect, a small pilot incision was made using a sterile Surgical Blade No. 11 (AD Surgical; Sunnyvale, CA), and the 5-gauge biopsy needle was inserted into the pilot incision ∼1 cm below the fascia. Approximately 50–80 mg of skeletal muscle was removed using a double chop method with applied suction. Following biopsies, tissue was rapidly teased of blood and connective tissue. A portion of the tissue (∼10–20 mg) was preserved in freezing media for histology (OCT; Tissue-Tek, Sakura Finetek Inc; Torrence, CA, USA), slowly frozen in liquid nitrogen-cooled isopentane, and subsequently stored at −80°C until histological analysis. Another portion of the tissue (∼30–50 mg) was placed in pre-labeled foils, flash frozen in liquid nitrogen, and subsequently stored at −80°C until other assays described below.

### Deep Proteomics for the Manual Interrogation of Denervation and Senescence-related Proteins

#### Sample Preparation

Sarcoplasmic protein isolation for all samples occurred at Auburn University using a previously described method [34], and the complete methods used have been previously described with a subset of the current participants [35]. Approximately 30 mg of tissue was homogenized using tight-fitting pestles in 500 µL of 25 mM Tris, pH 7.2, 0.5% Triton X-100 (with added protease inhibitors; Promega, Cat. No. G6521; Madison, WI, USA). Samples were then centrifuged at 1500 g for 10 min at 4°C and the soluble/sarcoplasmic fraction supernatants were collected. Due to resource constraints, 18 MA participants’ PRE and POST trained leg specimens and 9 younger participant specimens were used for further analysis.

Proteomics analysis of the sarcoplasmic fraction was performed at Seer, Inc. (Redwood City, CA, USA). For each sample, 250 μL soluble/sarcoplasmic fraction supernatant was subjected to the Seer Proteograph Assay protocol [36]. After the samples were loaded onto the SP100 Automation Instrument, they were subjected to protein corona formation and processing to generate desalted purified peptides protein identification using Reversed Phase (RP) Liquid Chromatography coupled to Mass spectrometry (LC-MS). To form the protein corona, Seer’s proprietary nanoparticles (NPs) were mixed with the samples and incubated at 37°C for 1 hr. Unbound proteins were removed prior to downstream wash, reduction, alkylation, and protein digestion steps which were performed according to Seer’s Proteograph Assay protocol [36].

#### LC-MS configuration

Peptides obtained from each of the five NP mixtures were separately reconstituted in a solution of 0.1% formic acid and 3% acetonitrile [37] spiked with 5 fmol μL PepCalMix from SCIEX (Framingham, MA, USA). Reconstitution volumes varied NPs to allow for constant peptide quantity for MS injection between samples regardless of starting volume (240 ng: NP1, 400 ng: NP2, 360 ng: NP3, 120 ng: NP4, and 320 ng: NP5). 4 µL of each sample were analyzed with a Ultimate3000 RLSCnano LC system coupled with a Orbitrap Fusion Lumos mass spectrometer (Thermo Fisher; Waltham, MA, USA). Peptides were loaded on an Acclaim PepMap 100 C18 (0.3 mm ID × 5 mm) trap column and then separated on a 50 cm μPAC analytical column (PharmaFluidics, Belgium) at a flow rate of 1 μL/min using a gradient of 5–25% solvent B (100% ACN) mixed into solvent A (100% water) over 26 min. The mass spectrometer was operated in Data Independent Acquisition mode using 10 m/z isolation windows from 380-1200 m/z and 3 s cycle time. MS1 scans were acquired at 60k resolution and MS2 at 30k resolution.

#### Data Processing

Data-independent acquisition LC-MS data were processed using Proteograph Analysis Suite (PAS) v2.1 (Seer, Inc) using the DIA-NN search engine (version 1.8.1) in library-free mode searching MS/MS spectra against an in silico predicted library based on Uniprot’s Homo Sapiens reference database (UP000005640_9606, download December 9^th^, 2022). Library free search parameters included trypsin, 1 missed cleavage, N-terminal Met excision, fixed modification of Cys carbamidomethylation, peptide length of 7-30 amino acids, precursor range of 300-1800 m/z, and fragment ion range of 200-1800 m/z. Heuristic protein inference was enabled, MS1 and MS2 mass accuracy was set to 10 ppm. Precursor FDR was set to 0.01, and PG q-value was set to 0.01. Quantification was performed on summed abundances of all unique peptides considering only precursors passing the q-value cutoff. PAS summarizes all nanoparticle values for a single protein into a single quantitative value. Specifically, a single protein may have been measured up to five times, once for each nanoparticle. To derive the single measurement value, PAS uses a maximum representation approach, whereby the single quantification value for a particular peptide or protein group represents the quantitation value of the NP which most frequently has measured any given proteins across all samples. The relative abundances of protein targets were obtained by normalizing raw spectral values for each identified protein to total spectra within-participant. After normalization, for proteins that were not detected, values were set at zero with the assumption the protein was in low abundance or not detected.

An *a priori* systematic search was performed to filter the number of proteins used for the current analysis. Specifically, Gene Ontology (GO; http://geneontology.org, [38]) and UniProt (https://www.uniprot.org/, [39]) were used to search terms “denervation”, “response to denervation involved in regulation of muscle adaptation” (GO:0014894), “neuromuscular junction” (GO:0031594), “neuromuscular junction development” (GO:0007528), “skeletal muscle innervation”, “axonogenesis involved in innervation” (GO:0060385), “skeletal muscle tissue regeneration” (GO:0043403), “axon regeneration” (GO:0031103), “aging”, “cellular senescence” (GO:0090398), “replicative senescence” (GO:0090399), “positive regulation of cellular senescence” (GO:2000774), or “senescent cells”. After the list was finalized, proteins belonging to these GO terms were manually interrogated. This approach yielded 65 proteins, with 46 being related to muscle-nerve communication and 19 being related to cellular senescence.

### SASP protein detection in muscle tissue

Sarcoplasmic protein isolates (80 µg protein) from the trained leg of a subset of MA participants (n=13) and younger participants (n=5) were subjected to a customized antibody-based fluorometric array containing SASP proteins (RayBiotech; Peachtree Corners, GA, USA; Cat. No.: AAH-CYT-G5-4). Samples were diluted 80 µg of total protein in 120 µl of blocking buffer and added into the array to incubate for two hours. After washing, the arrays were incubated first, with a biotin-conjugated anti-cytokine antibody mix and then a 555 streptavidin-fluorophore, both for two hours. All incubations were performed at room temperature with gentle rotation and washes before the next step. The array was then read using an MDC GenePix 4200A Microarray Scanner (Molecular Devices; San Jose, CA, USA). Expression readings for each SASP marker were calculated as follows (Equation 1):

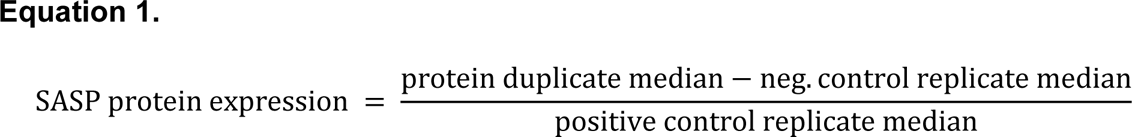

Zeros were substituted for proteins that yielded negative values (i.e., the signal was less than the negative control). All data are presented as relative expression units.

### Immunohistochemistry

Due to sample quality issues, n=20 MA participants (55±8 years old, n=15 females, n=5 males) and n=11 Y participants (23±4 years old, n=6 females, n=5 males) were used for immunohistochemistry. OCT-preserved samples were sectioned at 7 µm thickness using a cryotome (Leica Biosystems; Buffalo Grove, IL, USA) and adhered to positively charged slides. Participants’ trained and untrained leg muscle samples were placed on the same slides, as well as the PRE and POST sections to avoid batch-to-batch variation. Sections were then stored at -80°C until various staining procedures described in the following paragraphs.

The following methods were employed for the detection of denervated myofibers, fiber type-specific cross-sectional areas (fCSA) and myonuclei quantification. Slides were air-dried for 90-120 minutes prior to a 10-min acetone incubation at -20°C. Slides were then incubated with 3% H_2_O_2_ for 10 minutes at room temperature followed by an incubation with autofluorescence quenching reagent for 1 min (TrueBlack; biotium, Fremont, CA, USA). Next, a block containing 5% goat serum, 2.5% horse serum, and 0.1% Triton-X was applied and incubated for 1 h at room temperature. Another block was applied with streptavidin and then biotin solutions (Streptavidin/Biotin Blocking Kit, Vector Labs; Newark, CA, USA; Cat. No.: SP2002) at room temperature for 15 min each. Following blocking, slides were incubated overnight at 4°C with a primary antibody cocktail of NCAM (mouse IgG1, 1:50, Developmental Studies Hybridoma Bank; Iowa City, IA, USA; Cat. No. 5.1H11), myosin heavy chain I (mouse IgG2b, 1:100, Developmental Studies Hybridoma Bank; Cat. No. BA-D5-c), dystrophin (rabbit IgM, 1:100, GeneTex; Irvine, CA, USA; Cat. No.: GTX59790), and 2.5% horse serum in PBS. The next day, sections were incubated for 90 min in secondary biotin solution (anti-mouse IgG1, 1:1000, Jackson ImmunoResearch; West Grove, PA, USA), followed by a 60-min incubation with a secondary cocktail containing AF647 anti-mouse IgG2b (1:200, Thermo Fisher Scientific; Cat. No.: A21242), AF555 anti-rabbit IgM (1:200, Thermo Fisher Scientific; Cat. No.: A21428), and Streptavidin, AF488 Conjugate (1:500, Thermo Fisher Scientific; Cat. No.: S32354). Lastly, slides were stained with 1:10,000 DAPI (4’,6-diamidino-2-phenylindole, Thermo Fisher Scientific; Cat. No.: D3571) for 10 min at room temperature before coverslips were applied with PBS and glycerol (1:1) as mounting medium. Between each step, slides were washed in 1x phosphate buffered saline (PBS).

For quantification of satellite cells (Pax7+) and p21+ cells, a similar protocol was used. However, following the streptavidin and biotin block, the slides were incubated overnight at 4°C with a primary antibody cocktail of Pax7 (mouse IgG1, 1:20, Developmental Studies Hybridoma Bank) and 2.5% horse serum in PBS. The following day, slides were incubated for 90 min in secondary biotin solution (anti-mouse IgG1, 1:1000, Jackson ImmunoResearch), followed by a 60-min incubation with secondary streptavidin (1:500, SA-HRP, Thermo Fisher Scientific; Cat. No.: S-911), and a 20-min incubation with TSA-555 (1:200, Thermo Fisher Scientific, Cat. No.: B-40957). Next, a block containing 5% goat serum, 2.5% horse serum, was applied, and incubated for 1 h at room temperature. Similar steps from the fCSA/myonuclei/NCAM stain were applied except that the primary antibody cocktail included p21 (rabbit IgG, 1:200, Abcam; Cambridge, MA, USA; Cat. No. ab109199), dystrophin (mouse IgG2b, 1:100, Developmental Studies Hybridoma Bank; Cat. No. Mandys8 (8H11), and 2.5% horse serum in PBS. The next day sections were incubated for 90 min in secondary biotin solution (anti-rabbit IgG, 1:1000, Jackson ImmunoResearch) followed by a 60-min incubation with a secondary cocktail containing AF647 anti-mouse IgG2b (1:200) and Streptavidin, AF488 Conjugate (1:500). Again, slides were stained with 1:10,000 DAPI for 10 min at room temperature before coverslips were applied with PBS and glycerol (1:1) as mounting medium.

The last IHC performed was for quantification of p16+cells. The staining protocol was identical to the satellite cell and p21 staining protocol except that the primary cocktail containing p21 (rabbit IgG, 1:200, Abcam; Cambridge, MA, USA, ab109199), was substituted for p16 (rabbit IgG, 1:200, Abcam; Cambridge, MA, USA, ab51243).

Immediately following the mounting procedure for each stain, slides were imaged using a fluorescent microscope (Nikon Instruments, Melville, NY, USA) with the 20x objective lens. Four to six fields of view were captured per participant time point. fCSA and myonuclear number were analyzed using open-sourced software MyoVision [40]. Satellite cells, p21+ cells, and p16+ cells were manually quantified using Nikon NIS elements software (Nikon Instruments) and reported as number per 100 fibers.

### Statistical analysis

Statistical analyses were performed using SPSS (Version 26; IBM SPSS Statistics Software, Chicago, IL, United States). Prior to analysis, normality was assessed on all dependent variables using the Shapiro–Wilk’s test. All comparisons between MA and Y participants were analyzed using independent samples t-tests (for normally distributed data) or Mann-Whitney U-tests (for non-normally distributed data). Dependent samples t-tests were used for proteomic and SASP protein data to assess if training altered targets in MA participants, and Wilcoxon signed-rank tests were used for non-normally distributed data. All other variables were analyzed using multi-factorial (within-within) two-way (leg*time) repeated measures ANOVAs (or Freidman’s tests for non-normally distributed data), and in the event a significant interaction was observed (p<0.05), the model was decomposed for within- and between-leg comparisons at PRE and POST using parametric or non-parametric post hoc tests. Statistical significance was established at p<0.05. Eta square (*η*^2^) effect sizes are also provided for certain between-group comparisons and interactions, and effect size ranges were classified as follows: <0.06 = small effect, 0.06–0.14 moderate effect, and >0.14 large effect. All data herein are presented in figures and tables as means ± standard deviation values, and figures (aside from proteomic and SASP data) also present individual respondent values.

## RESULTS

### Participant characteristics

Baseline participant characteristics can be found in Table 2. Briefly, 26 participants (55±8 years old, n=17 females, n=9 males) completed the eight weeks of unilateral knee extensor resistance training, and general phenotype and training outcomes were collected for all these participants. However, as previously stated, only 20 MA participants yielded muscle samples suitable for histology (55±8 years old, n=15 females, n=5 males), and deep proteomics was performed for only 18 MA participants’ PRE- and POST-intervention trained leg due to resource constraints (55±8 years old, n=14 females, n=4 males). A subset of the fifteen Y participants (22±4 years old, n=9 females, n=6 males) were included for age comparison purposes. While estradiol levels in the Y females was numerically greater than the MA females, this was non-significant (502 ± 95 pg/mL versus 539 ± 119 pg/mL, p=0.465). This trend held with the removal of the four MA females on menopausal treatment prescriptions (p=0.155). Eleven of these 15 Y participants (23±4 years old, n=6 females, n=5 males) yielded tissue adequate for IHC comparisons to the 20 MA participants, and muscle tissue from 9 of these 15 Y participants (22±2 years old, n=7 females, n=2 males) were used for proteomic comparisons to the MA participants. Besides age (p<0.001), there were no significant differences in general anthropometrics between the 26 MA and 15 Y participants.

**Table 2.**
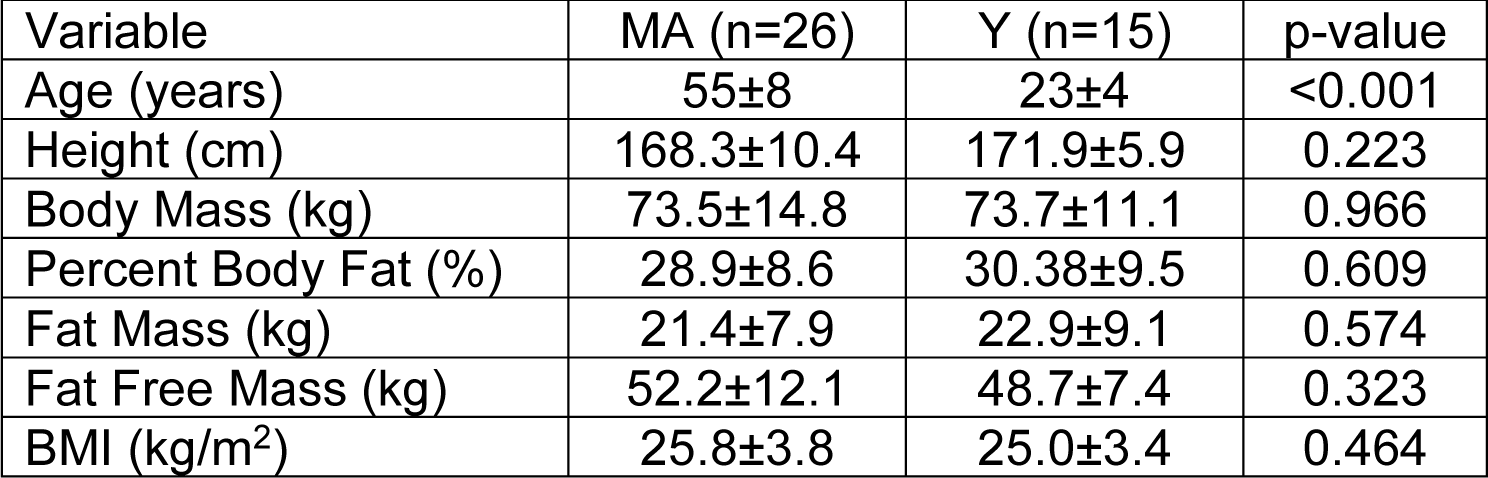
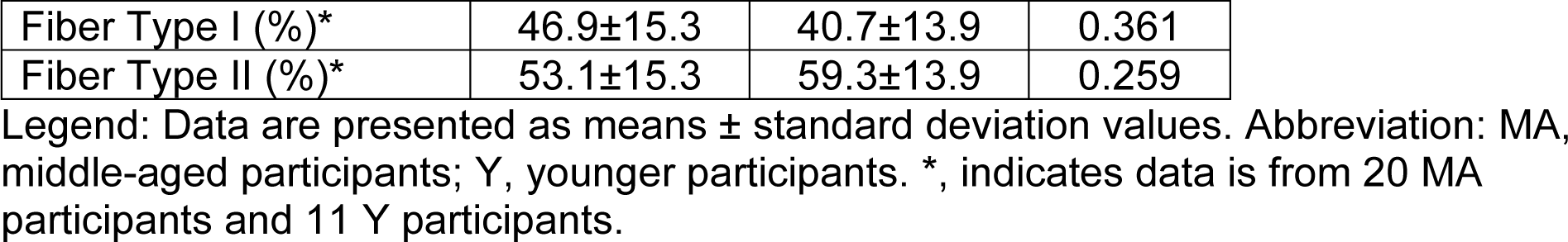
Participant characteristics.

### General Training Adaptations

Figure 1 depicts PRE and POST data in MA participants and basal state data in Y participants for VL mCSA, leg extension peak torque, and VL fCSA. While there was a significant leg (L) (p=0.047, ƞ^2^=0.174) and time (T) effect (p=0.048, ƞ^2^=0.067), there was not a significant leg x time (LxT) interaction for VL mCSA (trained leg: 20.19±5.20 cm^2^ to 20.64±5.03 cm^2^, untrained leg: 19.62±5.03cm^2^ to 19.62±5.05cm^2^, p=0.082, ƞ^2^=0.116, Fig. 1a). Additionally, there were no significant differences in values between MA and Y participants at any time point. A Friedman test was conducted to assess differences in leg extension peak torque across the groups. The test yielded a significant result (p=0.005), and Wilcoxon tests indicated significantly greater values in the trained leg after training (p<0.001) as well as the trained versus untrained leg at the post-intervention time point (p=0.011). However, there was no significant difference between MA and Y participants at any time point. Next, there were no significant effects of L, T, or LxT in type I (L: p=0.101; T: p=0.328; LxT: p=0.906, Fig. 1e), type II (L: p=0.923; T: p=0.195; LxT: p=0.437, Fig. 1f), or mixed fCSA (L: p=0.403; T: p=0.344; LxT: p=0.549, Fig. 1g). While type I fCSA yielded no significant differences between MA and Y participants at any time point, type II fCSA was significantly lower at all times in MA versus Y participants (p=0.005-0.017), and mixed fCSA was significantly lower only in the POST trained leg (p=0.035). Lastly, there were no significant effects of L, T, or LxT for average myonuclei for type I fibers (L: p=0.461; T: p=344; LxT: p=0.563) or type II fibers (L: p=0.451; T: p=0.420; LxT: p=0.209) (*data not graphed*). In addition, MA compared to Y participants had significantly more myonuclei per type I fibers in both legs at PRE (p=0.035 for both) but not at POST, or in type II fibers at PRE or POST (*data not graphed*).

**Figure 1.**
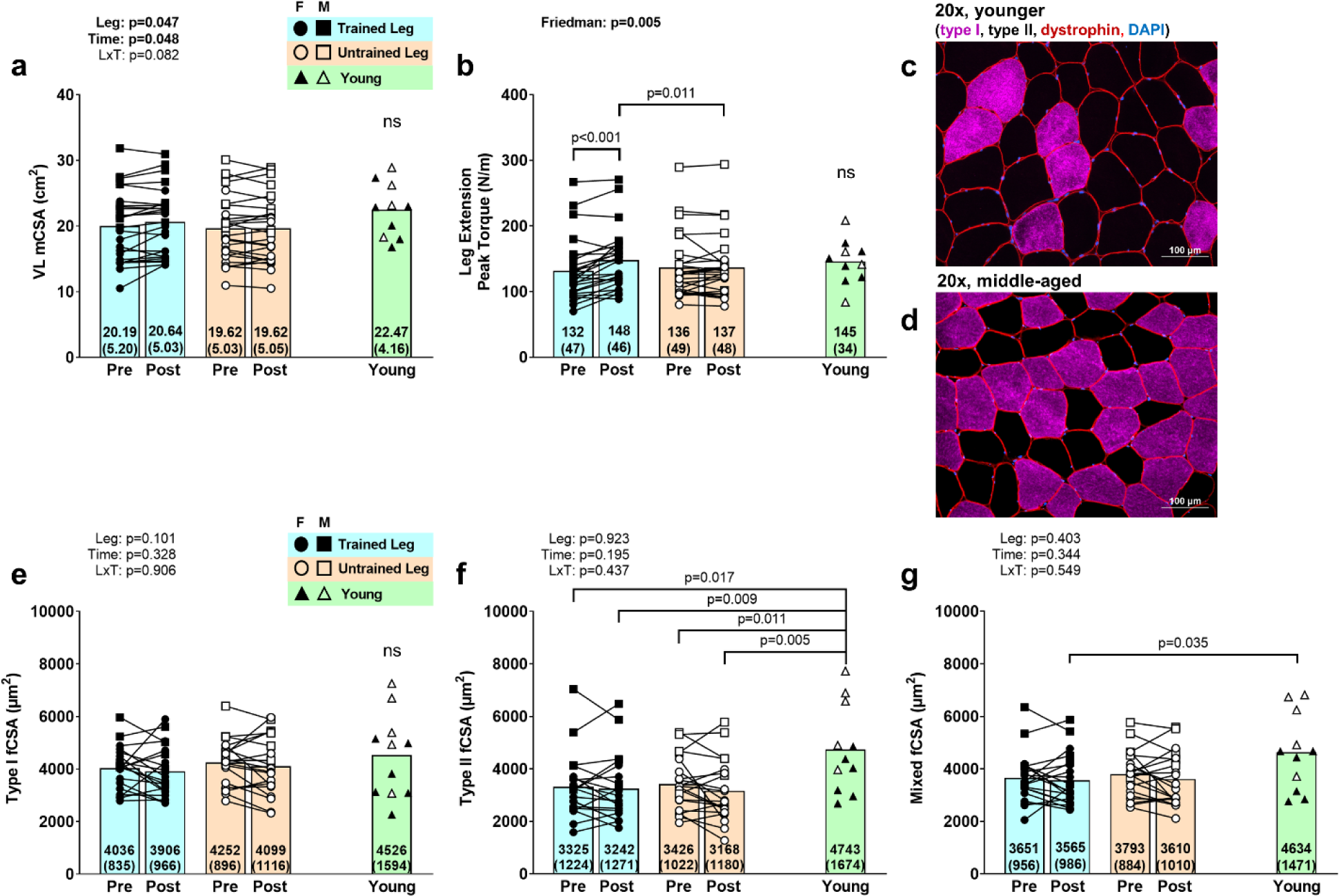
General training adaptations. Legend: Variables presented include vastus lateralis muscle cross-sectional area (a), leg extensor peak torque at 60°/sec in newton-meters (b), type I fiber cross-sectional area (e), type II fiber cross-sectional area (f), and mixed fiber cross-sectional area (g). Panels (a) and (b) contain n=26 middle-aged (MA) participants (55±8 years old, n=17 females, n=9 males) and n=10 Y participants (23±4 years old, n=6 females, n=4 males). Panels (e-g) contain n=20 MA participants (55±8 years old, n=15 females, n=5 males) and n=11 Y participants (23±4 years old, n=6 females, n=5 males). Representative images for fiber cross-sectional area for Y and MA participants are presented in panels c and d. Data are presented as mean values for PRE and POST intervention MA-trained and MA-untrained leg, and the Y comparator group. Individual responses are also illustrated, with open circles and squares indicating females and males, respectively, for the MA-untrained leg and closed circles and squares indicating females and males, respectively, for the MA-trained leg. Open and closed triangles indicate Y females and males, respectively.

### Denervated myofibers

For NCAM+ fiber quantification, a total of 184±78 and 162±67 fibers were counted prior to the eight weeks and 187±88 and 183±67 fibers after eight weeks for the trained and untrained legs, respectively. A Friedman test indicated that the percent of NCAM+ fibers in MA participants was not significantly different between legs for mixed (p=0.180, Fig. 2a), type I (p=0.061, Fig. 2b), or type II fibers (p=0.865, Fig. 2c). Interestingly, for both mixed NCAM+ fibers and NCAM+ type I fibers, the percentage of PRE trained (p≤0.006) and POST untrained leg (p≤0.010) were significantly higher than the Y cohort according to Mann-Whitney U tests. In contrast, only the POST trained leg was significantly higher than the Y cohort for percentage of type II NCAM+ fibers (p=0.022) according to a Mann-Whitney U test.

**Figure 2.**
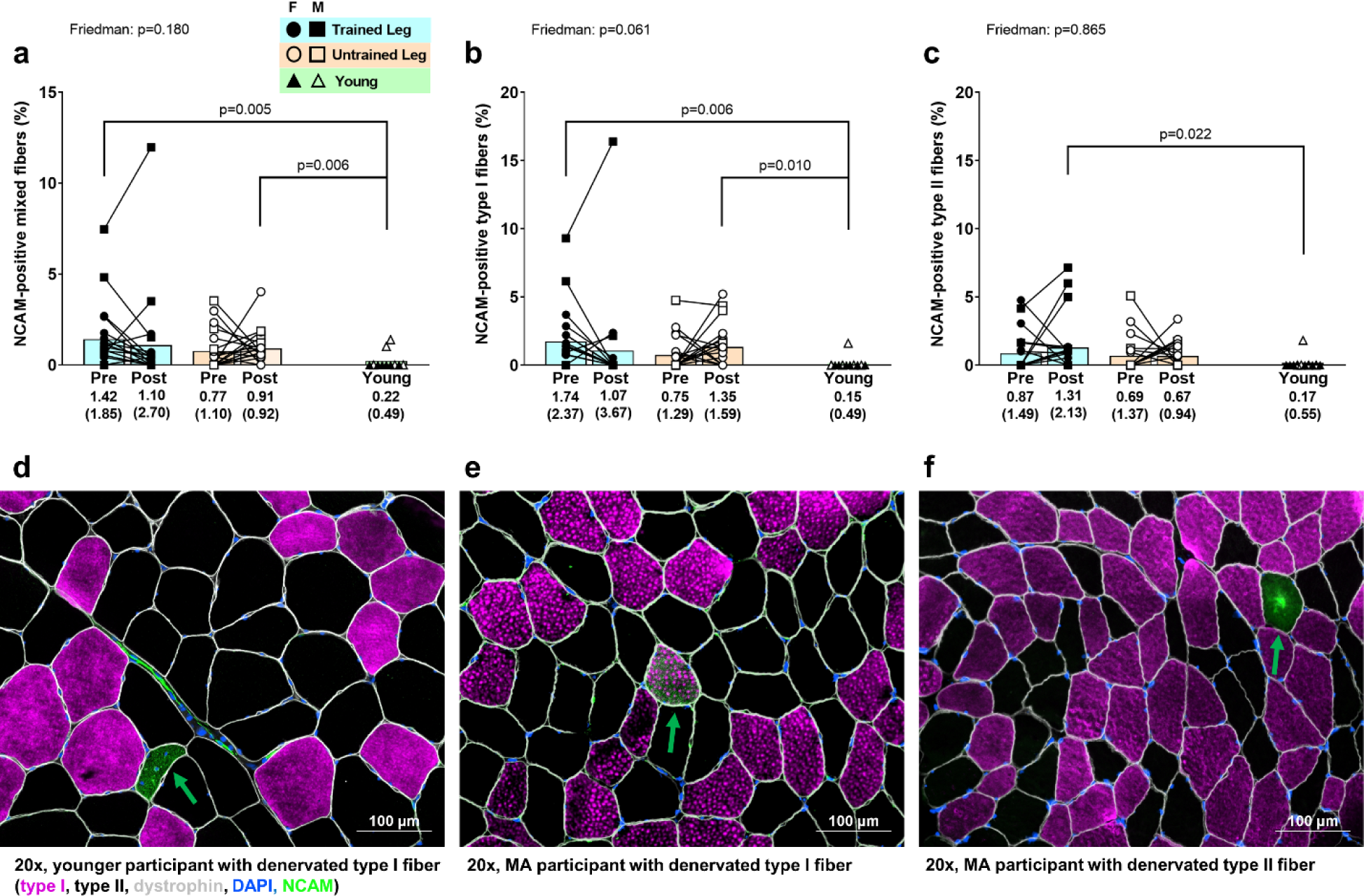
Immunohistochemistry for denervated myofibers. Legend: Variables presented include percent of mixed fibers that were NCAM+ (a), percent of type I fibers that were NCAM+ (b), percent of type II fibers that were NCAM+ (c). Panels a-c contain n=20 middle-aged (MA) participants (55±8 years old, n=15 females, n=5 males) and n=11 Y participants (23±4 years old, n=6 females, n=5 males). Representative 20x images are presented in panels d-f whereby green arrows depict NCAM+ myofibers. Data are presented as mean values for PRE and POST MA-trained and MA-untrained leg, and the Y comparator group. Individual responses are also illustrated, with open circles and squares indicating females and males, respectively, for the MA-untrained leg and closed circles and squares indicating females and males, respectively, for the MA-trained leg. Open and closed triangles indicate Y females and males, respectively.

### Satellite Cells and Senescent Cells

A significant LxT interaction was evident for satellite cells (trained leg: 7.9±3.4 to 10.4±6.0 cells/100 fibers; untrained leg: 8.5±4.2 to 8.2±1.8 cells/100 fibers, p=0.037, ƞ^2^=0.209, Fig. 3a). While PRE to POST values significantly increased in the training leg (p=0.020), POST values between legs were not significantly different (p=0.091). Additionally, satellite cell number for both MA participant legs at PRE and the untrained leg at POST were significantly lower (p≤0.018) than the Y cohort, but the values in the POST training leg were not (p=0.073). For p21+ cell quantification, a total of 206±79 and 189±69 fibers were counted prior to the eight weeks and 201±70 and 210±82 fibers after eight weeks for the trained and untrained legs, respectively. The Friedman test indicated the total number of p21+ cells in MA participants (both Pax7+ and Pax7-) was not significant between legs (p=0.138, Fig. 3b). However, compared to the Y cohort, Mann-Whitney U tests indicated that there was a significantly greater number of these cells in both legs, at both timepoints (p≤0.021). For p21+ satellite cells (p21+/Pax7+) in MA participants, a Friedman test revealed there was no significant effect between legs (p=0.162, Fig. 3e). Again, compared to the Y cohort, there was a significantly greater amount of these cells in both legs, at both timepoints (p≤0.036). For p16+ cell quantification, a total of 187±55 and 178±56 fibers were counted prior to the eight weeks and 204±55 and 203±69 fibers after eight weeks for the trained and untrained legs, respectively. The Friedman test was conducted for the total number of p16+ cells in MA participants and showed that there was a significant effect between legs (p=0.002, Fig. 3c). A Wilcoxon test indicated that these cells were significantly greater after training compared to the pre-training values in the trained leg (p=0.035). A Friedman test was also conducted to assess p16+ satellite cells (p16+/Pax7+) across legs in MA participants. The test yielded a significant result (trained leg: 0.42±0.91 to 1.07±1.34 cells/100 fibers; untrained leg: 0.90±1.28 to 1.47±1.55 cells/100 fibers, p<0.001, Fig. 3f), and a Wilcoxon test indicated that these cells were significantly greater after training compared to the pre-training values in the trained leg (p=0.015). At no time or leg was there a significant difference between MA and Y participants for this phenotype.

**Figure 3.**
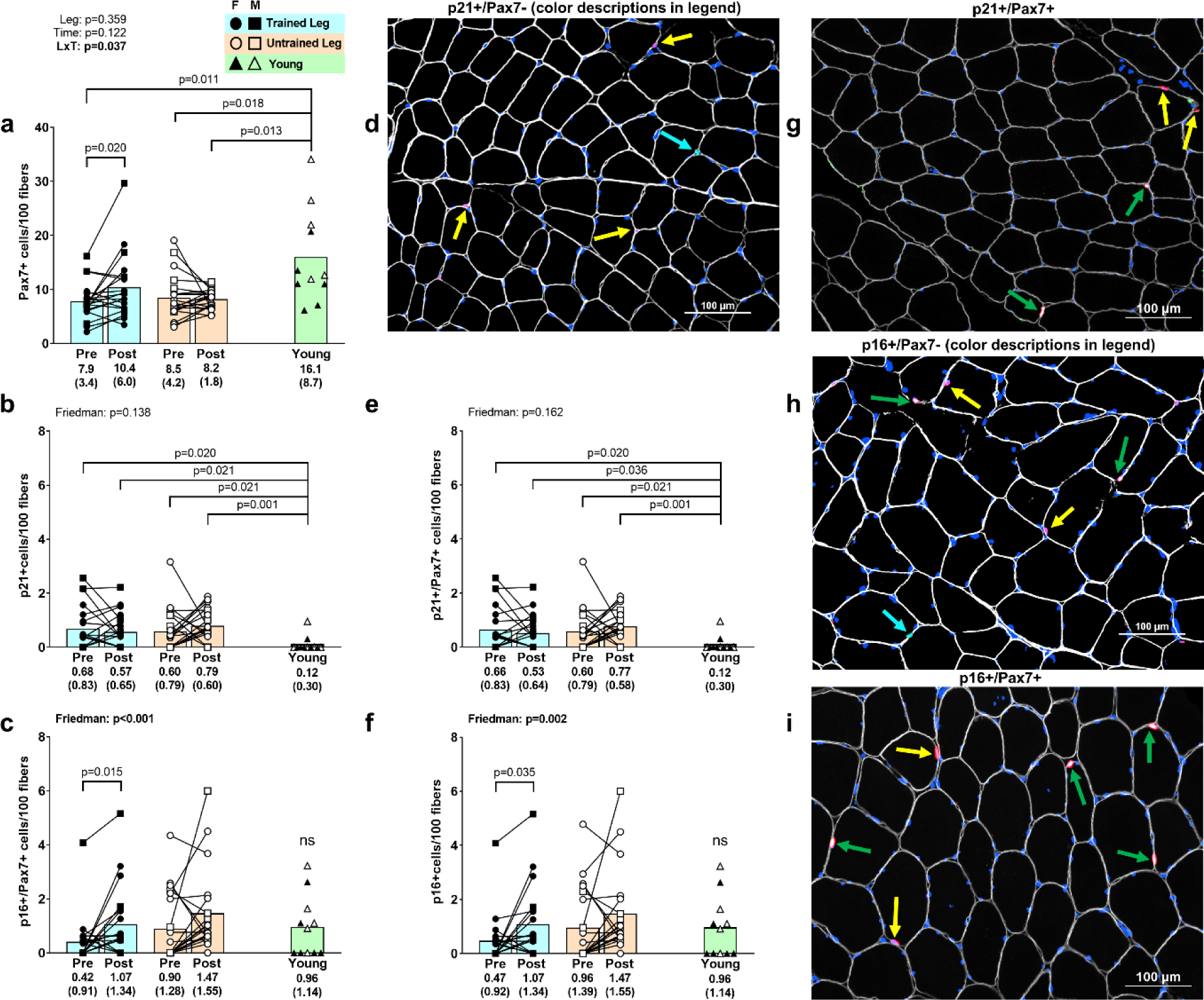
Satellite cells and senescent cells. Legend: Variables presented include Pax7+ cells per 100 fibers (a), p21+ cells/100 fibers (b), p21+/Pax7+ (satellite) cells/100 fibers (e), p16+ cells/100 fibers (c), and p16+/Pax7+ (satellite) cells/100 fibers. Graphs contain n=20 middle-aged (MA) participants (55±8 years old, n=15 females, n=5 males) and n=11 Y participants (23±4 years old, n=6 females, n=5 males). Data are presented as mean values for PRE and POST MA-trained and MA-untrained leg, and the Y comparator group. Individual responses are also illustrated, with open circles and squares indicating females and males, respectively, for the MA-untrained leg and closed circles and squares indicating females and males, respectively, for the MA-trained leg. Open and closed triangles indicate Y females and males, respectively. Representative images are presented in panels d, and g-i. Target colors in images are as follows: dystrophin (gray, pseudo color), Pax7 (red), DAPI (blue), and p21 or p16 (green). Arrows in images are as follows: p21-/Pax7+cells (yellow arrow), p21+ and Pax7-cells (cyan arrow), and p21+/Pax7+ cells (green arrow) in panels d and g, and p16-/Pax7+cells (yellow arrow), p16+ and Pax7-cells (cyan arrow), and p16+/Pax7+ cells (green arrow) in panels h and i.

Given the high number of p16+ and p21+ cells that co-localized with Pax7, we opted to provide a depiction of the percentage of senescent cells that were satellite cells (and vice versa) for Y and MA participants (trained leg only) (Fig. 4). Interestingly, while >90% satellite cells were not p16+ or p21+, most of the p16+ and p21+ cells co-localized with Pax7 (>90% on average in both MA time points and Y) indicating that most of the senescent cells were satellite cells in both age cohorts.

**Figure 4.**
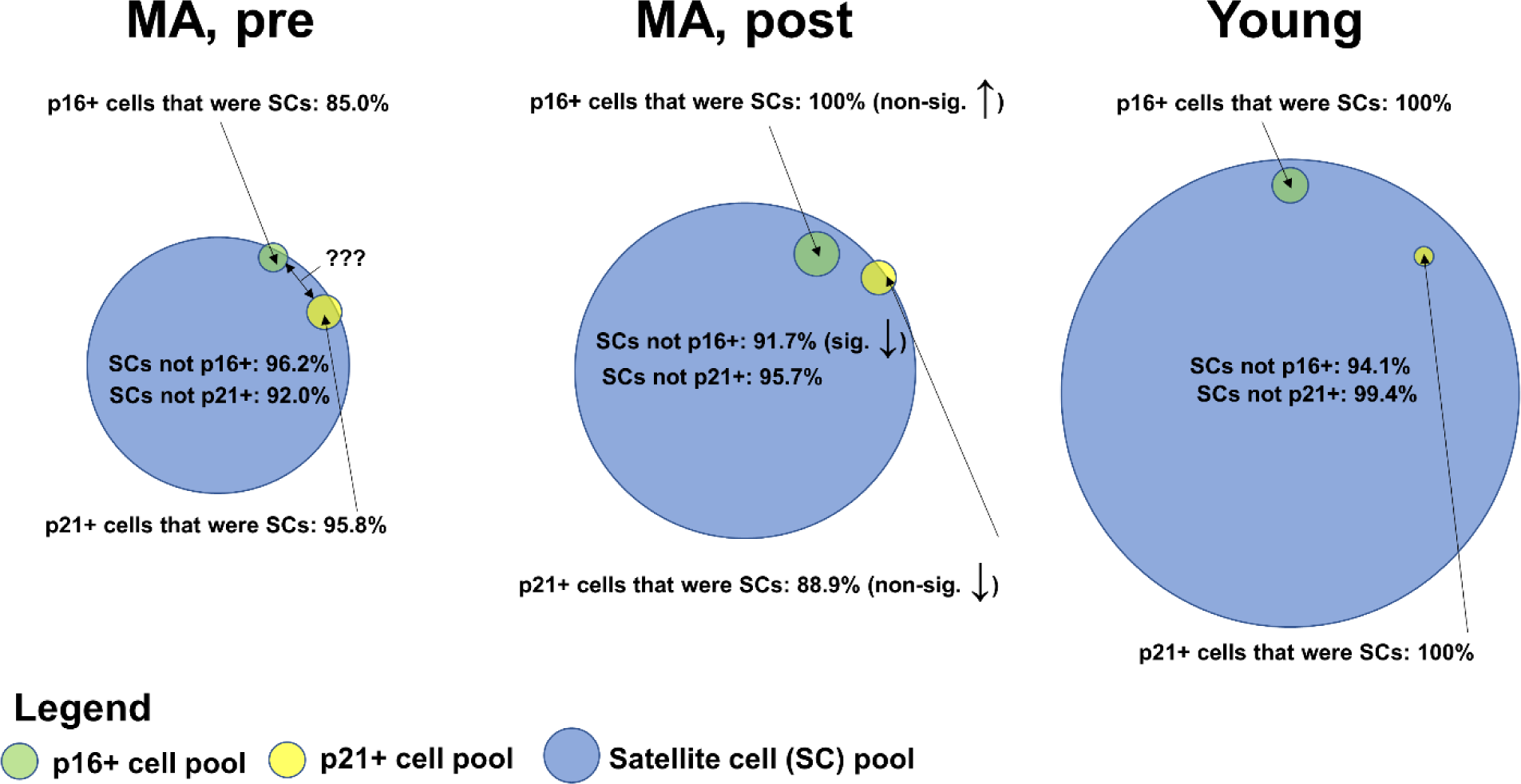
Percentage of satellite cells that were senescent cells and vice versa. Legend: Figure contains average cell percentages from the trained leg of n=20 middle-aged (MA) participants (55±8 years old, n=15 females, n=5 males) and n=11 Y participants (23±4 years old, n=6 females, n=5 males). Note: “???”, indicates that the percentage of p16+ and p21+ cell overlap is unknown given that this stain was not performed due to the limited number of filters utilized.

### Potential Sex Differences

While the number of male participants was small, we did perform exploratory *post hoc* three-way repeated measures ANOVAs on VL mCSA, leg extensor peak torque and histology outcomes (NCAM+ fibers, fCSAs, satellite cells, p16+/p21+ cells) to examine whether any sex-specific differences existed. The only significant interaction was a leg-by-time-by-sex interaction for percent type II NCAM+ myofibers in the trained leg only whereby PRE to POST change scores were lower in MA females versus males (females: -0.32±1.38%, males: 2.71±2.88%; p=0.014). Notably, these comparisons were not performed for proteomics, or the SASP protein array data given that data from fewer than 5 males were available for those variables.

### Changes in sarcoplasmic proteins related to muscle-nerve communication

The sarcoplasmic proteins related to muscle-nerve interaction are shown in Table 3. Of the 47 proteins, 13 significantly changed with training (all upregulated), 18 were significantly different prior to training compared to the Y cohort, and 20 were significantly different after training compared to the Y cohort.

**Table 3.**
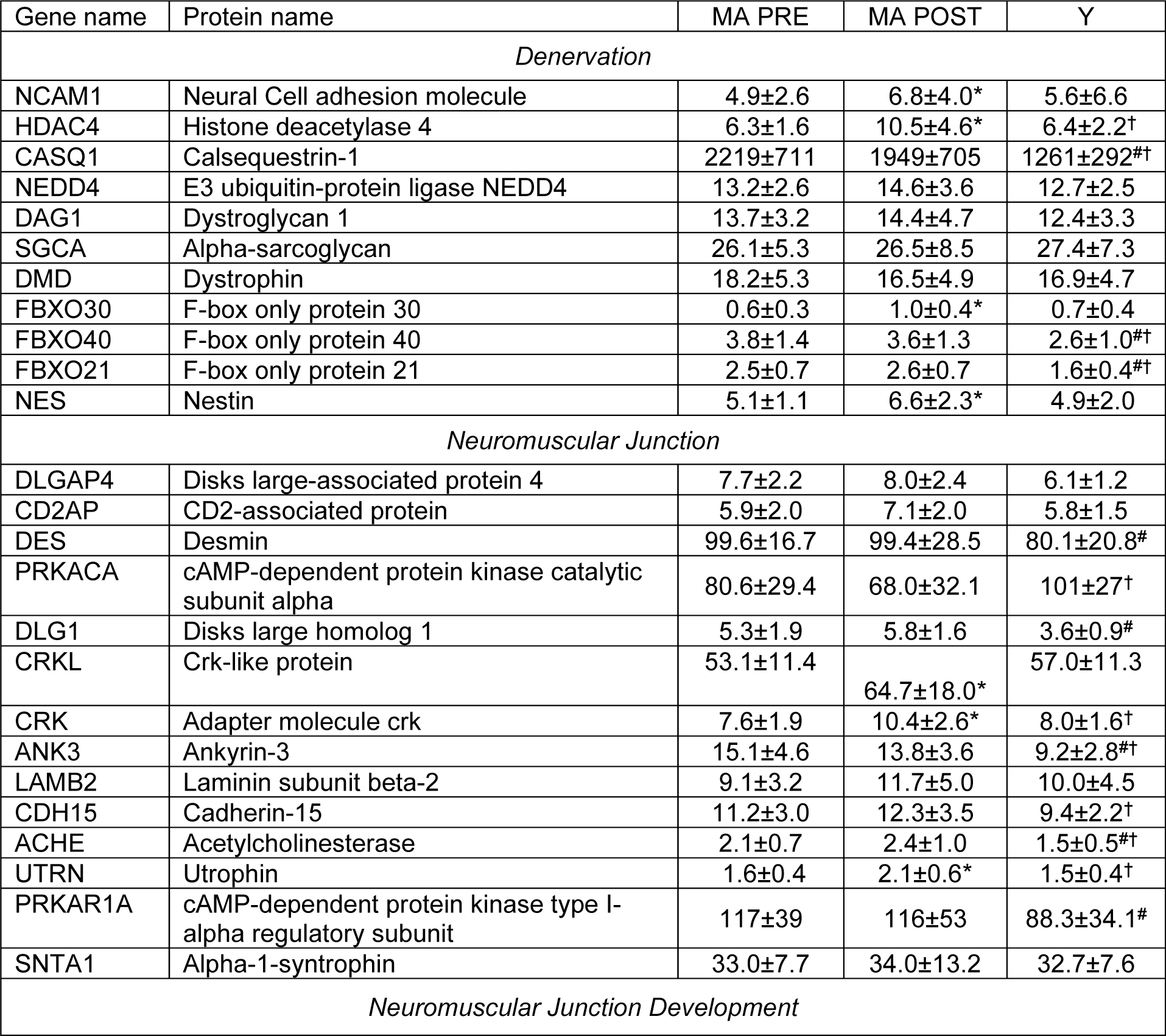
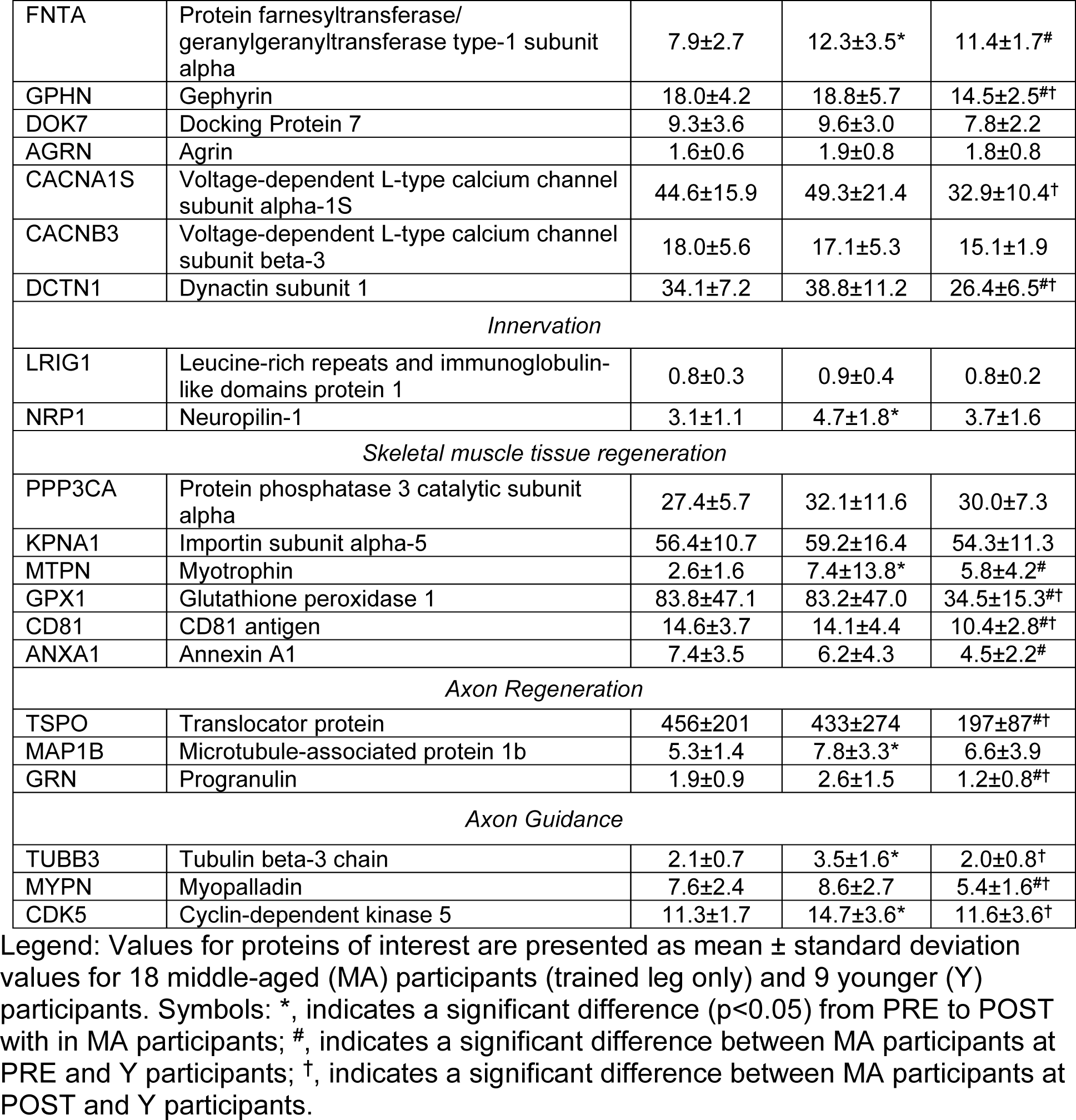
Expression of sarcoplasmic proteins related to muscle-nerve communication.

### Changes in sarcoplasmic proteins related to cellular senescence

There were 15 proteins of interest related to “cellular senescence” that were identified (Fig. 5). Eight of the 15 proteins were significantly altered with training (MAP2K1: 38.5±11.6 to 49.7±20.7, p=0.048; MAP2K3: 68.7±15.4 to 94.7±32.3, p=0.006; MAP2K4: 11.3±2.0 to 13.8±3.9, p=0.031; MAP2K6: 33.6± 8.1 to 51.3±25.5, p=0.012; MIF: 27.9±11.0 to 18.1±7.4, p=0.004; GLB1: 1.6±0.5 to 2.4±1.3, p=0.006; DNAJA3: 14.3±6.8 to 17.5±6.7, p=0.026; PML: 3.4±0.8 to 4.0±1.2, p=0.034). Prior to training, 6 of the 15 proteins were significantly different in MA versus Y participants (MAP2K1: 38.5±11.6 versus 62.3±18.1, p<0.001; MAPK14: 20.5±5.4 versus 30.4±7.8, p<0.001; NPM1: 36.0±15.9 versus 14.8±7.3, p<0.001; NUP62: 5.2±2.5 versus 3.2±0.9, p=0.014; DNAJA3: 14.3±6.8 versus 9.4±3.9, p=0.040). After training, 6 of the 15 proteins were significantly in MA versus Y participants (MAP2K6: 51.3±25.5 versus 32.4±7.5, p=0.035; NPM1: 39.2±18.3 versus 14.8±7.3, p<0.001; NUP62: 5.2±1.4 versus 3.2±0.9, p=0.001; CALR: 137±35 versus 113±18, p=0.026; DNAJA3: 17.5±6.7 versus 9.4±3.9, p=0.003; PML: 4.0±1.2 versus 3.1±0.5, p=0.025). Lastly, two search terms “Replicative senescence” and “Positive regulation of cellular senescence”, were not included in Figure 4, but are shown in Table 3 below.

To further interrogate whether cellular senescence was upregulated with training, we entered the seven proteins from the “cellular senescence” pathway that were upregulated, into PANTHER pathway analysis (MAP2K1, MAP2K3, MAP2K4, MAP2K6, GLB1, DNAJA3, PML) [41] and performed the PANTHER Overrepresentation Test whereby Fisher’s Exact and False Discovery Rate was calculated [42]. The test indicated that “cellular senescence” was predicted to be >100-fold upregulated (raw p-value = 5.99 x 10^-15^, FDR = 9.35 x 10^-11^).

**Figure 5.**
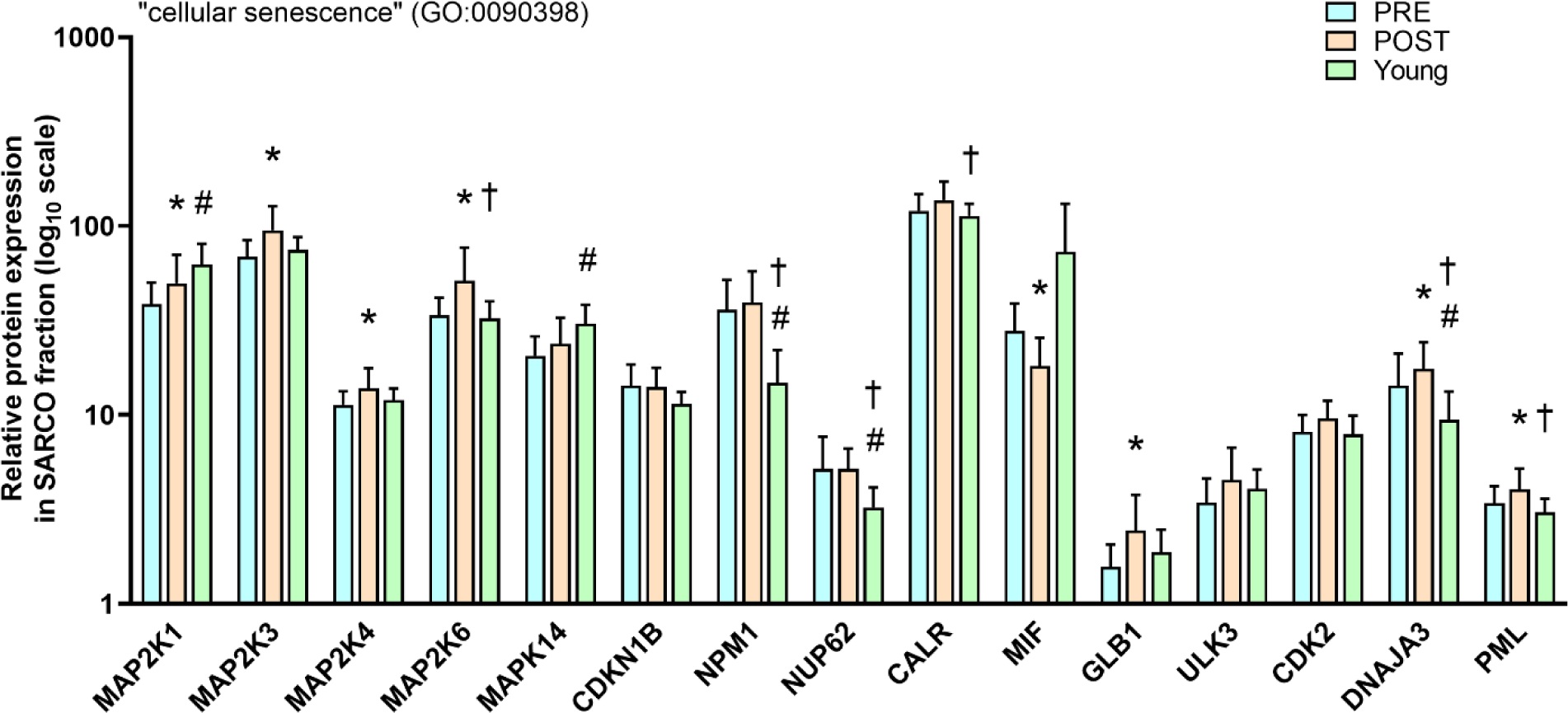
Expression of sarcoplasmic proteins related to cellular senescence. Legend: Data presented as mean ± standard deviation values from n=18 middle-aged (MA) participants (55±8 years old, n=14 females, n=4 males) and n=9 Y participants (22±2 years old, n=7 females, n=2 males). Data are scaled as log_10_ values to improve visualization. Symbols: *, indicates a significant difference (p<0.05) from PRE to POST in MA participants; ^#^, indicates a significant difference between MA participants at PRE and Y participants; ^†^, indicates a significant difference between MA participants at POST and Y participants.

**Table 4.**
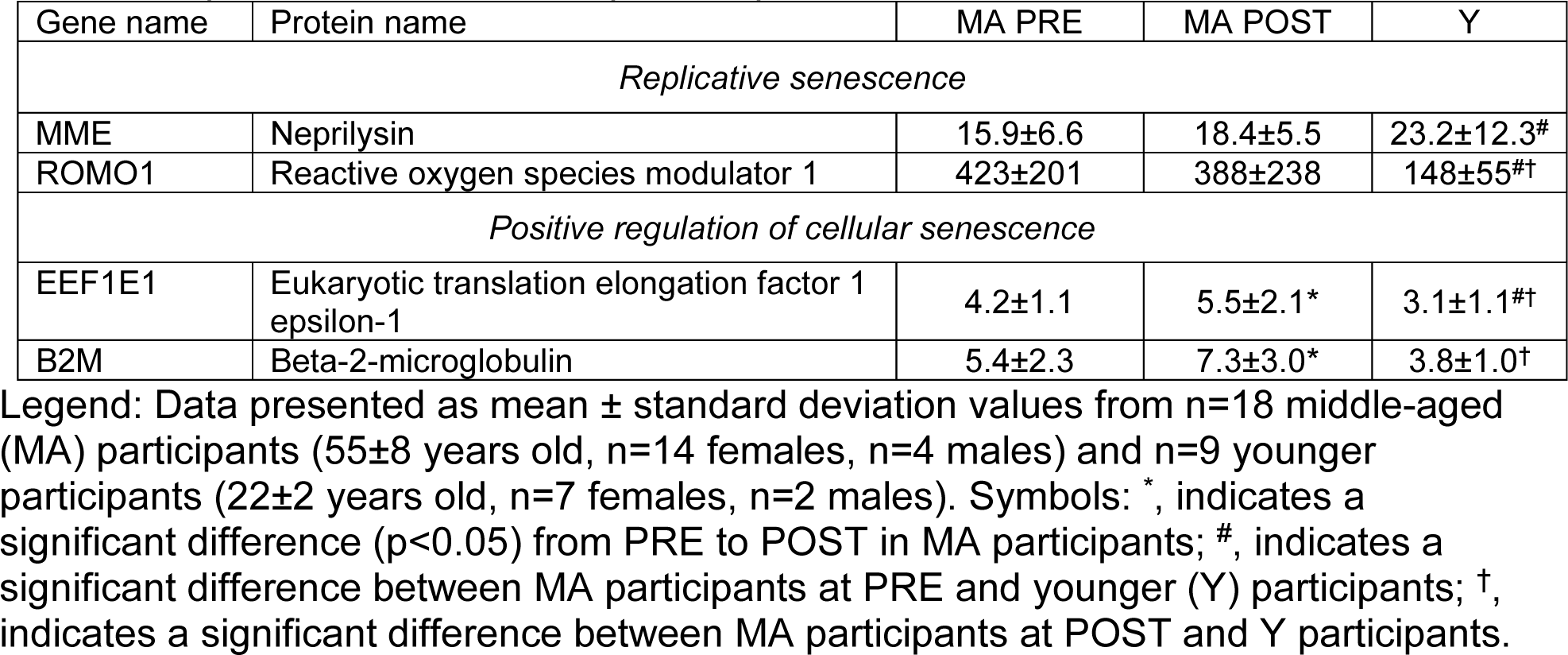
Expression of other sarcoplasmic proteins related cellular senescence.

### Changes in SASP protein expression

Based on comprehensive reviews [43, 44], 17 of the 160 targets provided by the antibody array were interrogated as SASP targets of interest (MCP-1, Osteopontin, IL-1α/-6/-8/-10, GM-CSF, GRO-a, IFN-g, MCSF, RANTES, HGF, IGFBP3/4, TGFβ1/2, and VEGF; Figure 6). One of the 17 SASP proteins were significantly altered with training in MA participants (IGFBP-3: 0.546±1.076 to 1.257±2.024, p=0.031). Prior to training, no SASP proteins were significantly different in MA versus Y participants. After training, two of the 17 SASP proteins were significantly different in MA versus Y participants (IGFBP-3: 1.257±2.024 versus 0.091±0.204, respectively; p=0.048; Osteopontin: 1.253±1.636 versus 0.185±0.258, respectively; p=0.025).

**Figure 6.**
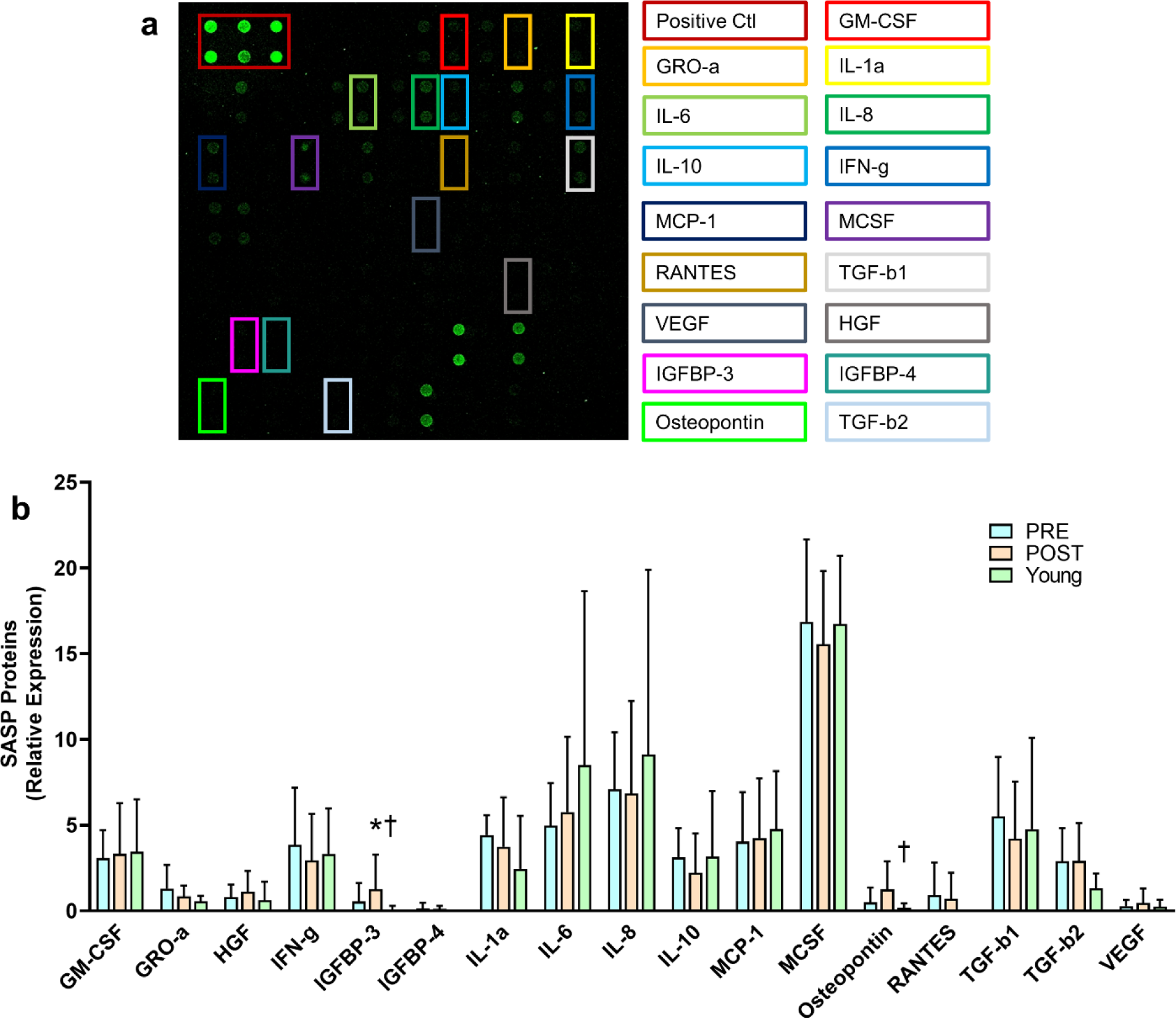
Expression of SASP proteins. Legend: (a) A representative array, where each sample was assayed in duplicate. The level of SASP protein expression is proportional to fluorescence intensity. SASP proteins are labeled and color matched on the right. (b) SASP protein expression data presented from n=13 MA participants (58±8 years old, n=3 males, n=10 females) and n=5 Y participants (22±1 years old, n=1 males, n=4 females). Symbols: *, indicates a significant difference (p<0.05) from PRE to POST in MA participants; ^†^, indicates a significant difference between MA participants at POST and Y participants.

## DISCUSSION

We sought to examine if denervated myofibers, senescent cells, and associated protein markers in MA individuals were affected with eight weeks of unilateral knee extensor resistance training, and whether training restores these markers to “youth-like” levels. Our primary findings are as follows: i) despite age-related differences in denervated myofiber content, resistance training did not affect this variable, ii) training did not alter p16+ and p21+ cell content or most SASP proteins, albeit training did increase several senescence-associated proteins, iii) training increased satellite cell content in the absence of myofiber hypertrophy, iv) there were more p21+ cells in MA compared with Y participants, and v) although most satellite cells were not senescent, most senescent cells were satellite cells. Finally, the applied and cellular responses seemed to be largely conserved between sexes, albeit limited sample size for males require more validation research in this regard.

The current literature is mixed as to whether exercise training can affect NCAM+ myofibers in older adults. Our null findings in the current study are congruent with other studies [15, 17], albeit one study has indicated that the area occupied by NCAM+ myofibers in muscle sections is reduced with resistance training [16]. Although we did not observe a training effect, it is interesting that the percentage of mixed and type I NCAM+ fibers were significantly greater at PRE in the trained leg of MA versus Y participants. Further, unlike PRE values, values of NCAM+ myofibers (both total and type I) in the trained leg at POST were not significantly different from the Y cohort. Also notable, several proteins related to muscle-nerve communication were upregulated in the trained leg of MA participants. We interpret these collective findings to suggest that, had the training intervention been longer or perhaps more intense, resistance training may have increased type I (but not type II) fiber innervation. This is a plausible hypothesis given that various reports indicate that active, MA individuals possess larger type I fibers relative to inactive counterparts while also possessing a greater degree of fiber type grouping, alluding to a greater degree of reinnervation [9, 45]. However, Soendenbroe et al. [17] recently reported that a training intervention that was twice as long in duration compared to the current study and had participants perform two additional leg exercises per session did not affect the number of NCAM+ fibers. Hence, it is currently unclear how shorter-term resistance training affects myofiber denervation, and more studies in both middle-aged and older participants are needed to draw more definitive conclusions.

Senescent cell number has been reported to be higher in aged skeletal muscle [25], and recent research interest has investigated how exercise affects this phenotype [46, 47]. While numerous studies have shown a decrease in senescent markers with exercise [21, 22, 48-50], our results suggest training does not affect p16+ or p21+ cells in skeletal muscle. There was an age effect with p21+ satellite cells whereby the MA participants had a significantly higher abundance compared to the younger cohort in both legs prior to and following the training intervention. Past reports have indicated that senescent cells are not altered or increase with mechanical overload [51-53]. However, it is noteworthy that studies examining p16+ or p21+ senescent cells with exercise have not indicated whether they are satellite cells. A key finding in our investigation is that most senescent cells co-localized with Pax7 in both age cohorts. Additionally, satellite cells increased in the trained leg, and this coincided with several senescent-related proteins increasing in the trained leg as well. Another interesting and related finding is that SASP proteins did not increase in the trained leg of MA participants despite several senescent proteins increasing, and SASP protein differences between MA and Y were marginal despite p21+ cells being higher in both legs of MA at all times. While speculative, we interpret this collective evidence to suggest that senescence signals in muscle tissue could primarily come from satellite cells, and that the increase in satellite cells observed with training drove several of the senescence-related protein (but not SASP protein) responses. This viewpoint is partially supported by a recent review [47] indicating that the expression of cyclin-dependent kinase inhibitors and other cellular senescence proteins could generally represent an increase in cell proliferation and differentiation rather than cellular senescence. This hypothesis may also, in part, explain why Dungan and colleagues reported a robust increase in p21+ cells 14 days following an acute resistance exercise bout in younger (21-39-year-old) participants [27]. Specifically, while the authors did not examine if these cells co-localized with Pax7, it remains possible that the increase in p21+ and SA β-Gal cells were a portion of the satellite cell pool that were primed for differentiation.

It is worth discussing the satellite cell findings in the context of other training adaptations observed in the present investigation. While there are disagreements in the literature about how aging affects satellite cells [54-59], the current data indicate MA individuals had fewer satellite cells than younger individuals prior to training. With training, however, MA participants were able to increase satellite cell numbers back to youth-like levels, albeit this was not accompanied with increases in fCSA or myonuclear number. Other studies have demonstrated similar increases in satellite cell content with resistance training in MA participants, but with mixed hypertrophic outcomes [60, 61]. It has been previously reported that satellite cell content prior to training (regardless of age) is linked to different hypertrophy outcomes, which could explain the lack of myofiber hypertrophy in the current study [62, 63]. In addition to the lack of fCSA increases with training, the lower number of satellite cells prior to training in MA versus younger participants could explain why fCSA, specifically type II fCSA, was lower in the MA participants. While we did not interrogate fiber-type specific satellite cell content, others have shown that older participants possess significantly less type II fiber-specific satellite cells which corresponds with smaller type II fCSA [61, 64, 65]. It should also not be discounted that, while myofiber hypertrophy was not observed in the trained leg of MA participants, the increase in satellite cells could have acted to facilitate other adaptations (e.g., connective tissue remodeling) as reported in murine models by Fry and colleagues [66, 67], and this could have acted to facilitate tissue-level hypertrophy. Notwithstanding, the current data supports that satellite cells increase with resistance training and, had training been longer or more rigorous, this may have eventually translated to an increase in myofiber hypertrophy.

A final topic worthy of discussion is the paradoxical findings between VL mCSA changes in MA participants’ trained leg (which trended upward) and the numerical decreases in fCSA values. We have observed this phenomenon on prior occasions with resistance training interventions [33, 68], and speculate that this is likely due to the limited number of myofibers sampled with histology. However, it is intriguing that fCSA values numerically decreased given that our past studies suggested that tissue-level and myofiber area increases, while not significantly associated, both directionally increase with resistance training. Although this is difficult to reconcile, we posit that this may be due to the mild nature of knee extensor-only resistance training. Alternatively stated, relatively modest training adaptations were observed, and it is likely that more robust adaptations would have been observed had participants performed multiple exercises targeting the knee extensor muscles (e.g., knee extensors, squats, deadlifts, etc.).

### Limitations

Our study was not without limitations. First, we pooled MA participants who consumed a nutritional supplement or placebo over the eight-week intervention. As stated, the resistance training arm of this protocol was designed to examine the current research question, and we had no reason to suspect that the outcome variables in the trained leg would be affected by the supplied supplementation. While we provided evidence of this in the methods section, readers should be aware of this study design limitation. It is also worth noting that proteomic data were obtained from tissue containing mixed myofibers and stromal cells. Single fiber (or single cell) isolation with downstream proteomics analysis for NCAM+ fiber-specific, or p16+/p21+ cell-specific protein expression profiles would have provided more insight as to how training affected the proteins of interest in a cell-specific fashion. Additionally, while we lacked tissue to perform additional p16/p21 IHC with other cell types may have been insightful (e.g., fibro-adipogenic cells). The training paradigm was isolated to a single leg, and results may have differed had multiple leg exercises or full body training been implemented. Having a younger cohort that trained would have also been beneficial to examine whether the training adaptations seen with the MA participants were similar in younger participants. Lastly, there were a limited number of male participants in the study which limited our power to determine whether sex-specific adaptations existed in MA participants.

## Conclusions

In summary, although eight weeks of unilateral leg resistance training did not affect denervated myofiber or p16+/p21+ senescent cell counts in MA individuals, several proteins associated with muscle-nerve communication and senescence were upregulated. We conclude that resistance training either needed to be more vigorous and/or longer in duration to eventually affect these outcomes or does not affect these outcomes. Hence, more research is needed in this regard to provide further clarity. We also provide continued support for the role of resistance training in increasing satellite cell numbers in MA participants. Finally, most p16+/p21+ senescent cells in the muscle tissue of younger and MA participants were Pax7+ satellite cells, and SASP proteins were generally not different between MA and Y participants. Thus, we posit that these signatures represent the existence of a subpopulation of satellite cells, rather than proinflammatory SASP-secreting cells, being the primary source of senescent-like cells in skeletal muscle. However, again, more insight is needed to validate this hypothesis.

## ACKNOWLEDGEMENTS

We thank the participants who volunteered and participated in the study. We thank Seer, Inc. and their team members (David Hill, Mara Riley, Ryan Hill, Aaron S. Gajadhar, and others) for assisting us through the various procedures needed to perform proteomics analysis including performing pilot feasibility experiments using the Proteograph Product Suite. Shao-Yung Chen is an employee of Seer, Inc. who was largely responsible for the execution of the pilot feasibility experiments, although he had no role in the study design. The study design and outcome variables of this pooled analysis sought to examine training outcomes unrelated to the nutritional supplement administered herein, and the study design was created by B.A.R. and M.D.R. to fulfill dissertation requirements for B.A.R. However, the reader should be aware that M.D.R. and T.N.Z. have published research examining the effects of the administered supplement on unrelated cellular metabolites and metabolic enzymes *in vitro* and in humans, and current exploratory analyses from blood specimens and tissue specimens from the untrained leg of MA participants are ongoing in this regard. None of the other co-authors have apparent conflicts of interest in relation to these data.

## DATA AVAILABILITY

Raw data related to the current study outcomes will be provided upon reasonable request by emailing the corresponding author.

## FUNDING INFORMATION

Funding for participant compensation, histology, and proteomics was made possible through gift and discretionary funds from the M.D.R. laboratory, discretionary funds from the A.D.F. laboratory, and indirect cost sharing (generated from various unrelated contracts) from the School of Kinesiology. M.C.M. was fully supported through a T32 NIH grant (T32GM141739), and D.L.P. was fully supported by a Presidential Graduate Research Fellowship funded by Auburn University’s President’s office, the College of Education, and School of Kinesiology.

